# Cardiac and Skeletal Actin Substrates Uniquely Tune Cardiac Myosin Strain-Dependent Mechanics

**DOI:** 10.1101/348920

**Authors:** Yihua Wang, Katalin Ajtai, Thomas P. Burghardt

## Abstract

Native cardiac ventricular myosin (βmys) translates actin under load by transducing ATP free energy into mechanical work on actin during muscle contraction. Unitary βmys translation of actin is the myosin step-size. In vitro and in vivo βmys regulates contractile force and velocity by remixing 3 different step-sizes with stepping frequencies autonomously adapted to workload. Cardiac and skeletal actin isoforms have a specific 1:4 stoichiometry in normal adult human ventriculum. Human adults with inheritable hypertrophic cardiomyopathy (HCM) up-regulate skeletal actin in ventriculum suggesting that increasing skeletal/cardiac actin stoichiometry also adapts myosin force-velocity to respond to the muscle’s inability to meet demand.

Nanometer scale displacement of quantum dot (Qdot) labeled actin under resistive load when impelled by βmys measures single myosin force-velocity in vitro in the Qdot assay. Unitary displacement classification constraints introduced here better separates myosin based signal from background upgrading step-size spatial resolution to the sub-nanometer range. Single βmys force-velocity for skeletal vs cardiac actin substrates was compared using the Qdot assay.

Two competing myosin strain-sensitive mechanisms regulate step-size choices dividing mechanical characteristics into low- and high-force regimes. The actin isoforms alter myosin strain-sensitive regulation such that onset of the high-force regime, where a short step-size is a large or major contributor, is offset to higher loads by a unique cardiac ELC N-terminus/cardiac-actin contact at Glu6/Ser358. It modifies βmys force-velocity by stabilizing the ELC N-terminus/cardiac-actin association. Uneven onset of the high-force regime for skeletal vs cardiac actin dynamically changes force-velocity characteristics as skeletal/cardiac actin fractional content increases in diseased muscle.

## INTRODUCTION

Cardiac myosin has a 140 kDa N-terminal globular head called subfragment 1 (S1) and an extended α-helical tail domain. Tail domains form dimers that self-assemble into myosin thick filaments with S1’s projecting outward from the core in a helical array *^1^*. Thick filaments interdigitate with actin thin filaments in striated muscle and slide relatively during contraction *^2^*. S1 has the ATP and actin binding sites (the motor) and a lever arm whose rotary movement cyclically applies tension to move a load when myosin is strongly actin bound *^3^*. Actin binding sites in the S1 heavy chain cover solvent exposed regions of the molecule including structured loops (C-loop *^4, 5^*, myopathy loop, and loop 3), unstructured loop 2 *^6^*, and other peptide fragments in the motor and within the 50 and 20 kDa tryptic fragments of the S1 heavy chain *^7–9^*.

The lever arm converts torque generated by the motor into linear displacement (step-size) and undergoes shear strain due to the resisting stress. Strain affects the lever arm and the lever arm bound essential and regulatory light chains (ELC and RLC). The ~20 kDa RLC stabilizes the lever arm *^10–12^* and disease implicated RLC mutants lower velocity, force, and strain sensitivity *^13^* suggesting they alter lever arm processing of shear stress *^14, 15^*. The ~25 kDa skeletal myosin ELC (A1) has an N-terminus extension containing sites for ELC/actin binding localized in the N-terminal fragment and proposed to involve actin sites near to residue D363 *^16^*. Subsequent work on atrial ELC localized the ELC/actin binding site to the N-terminal 11 residues *^17^*. A structural model for S1 bound F-actin built from skeletal S1 and skeletal actin crystal structures indicated a close match for lysine residues K3 and K4 on the ELC A1 with actin’s E361 and E364 *^18^*. This ELC/actin interaction also has residue V6 on the ELC A1 and T358 on skeletal actin proximal and with side chain Cγ atoms separated by ~3.9 Å.

Skeletal and cardiac actin isoforms have a highly conserved sequence with >98% sequence identity. Their sequences differ at positions 2 and 3 where Glu-Asp is reversed to Asp-Glu in skeletal and cardiac isoforms and at the sites Leu299Met and Thr358Ser for Leu and Thr in the skeletal actin. Various actin isoforms share the same conformation *^19^* likely assuring near identical three-dimensional structures for skeletal and cardiac actin. A S1 bound F-actin homology model using cardiac ELC and actin sequences, and based on the skeletal sequences model *^18^*, places the S358T actin modification in a hydrogen bond with E6 in the cardiac ELC. We propose that this interaction specific to cardiac muscle modifies the cardiac motor force-velocity characteristics detected by single myosin mechanics.

Qdot labeled actin in vitro motility assay (Qdot assay) has myosin immobilized on a planar glass substrate impelling single Qdot labeled F-actin with nanometer scale actin displacement measured using super-resolution*^20, 21^* and total internal reflection fluorescence (TIRF) microscopy *^22^*. The assay has negligible compliance faithfully characterizing single myosin motor displacement of actin *^23–25^*. We found using the unloaded Qdot assay that ventricular cardiac myosin ELC N-terminal extension binds skeletal actin to modulate myosin step-size *^23^*. We demonstrated that βmys in vitro has three distinct unitary step-sizes of ~8, 5, and 3 nm moving unloaded actin and suggested that step-size selection depends on strain dependent regulation. In this model, 8 and 5 nm steps are complete transduction cycles while the 3 nm step, sometimes observed in the Qdot assay as a separate step, is coupled to a preceding 5 nm step. Strain inhibited ADP release *^26^* sometimes delays the 3 nm portion of some lever arm swings enough to be observed separately. Necessary generalization of this model to accommodate in vivo data and loaded Qdot assay data, as described below, caused us to define strain-regulated myosin/actin displacement pathway fluxes (rather than overall step-size probabilities) for particular step-sizes or step-size combinations. The pathways identified by the unloaded Qdot assay are now designated 8, 5, and 5+3 nm step-size pathways.

The complexity and density of the muscle sarcomere prohibited sensing single myosins in vivo until the visible light transparent zebrafish embryo opened a window into time-resolved single myosin mechanics. We imaged rotation of a single myosin lever arm tagged with photoactivatable green fluorescent protein in transgenic zebrafish embryo skeletal and cardiac muscle as it converted motor generated torque into unitary step displacement *^27, 28^*. Single cardiac myosin imaging of zebrafish embryo hearts suggested a 4 pathway network generating the original 8, 5, and 5+3 nm step-sizes plus a solo-3 nm step-size. The in vivo system added the context of the intact auxotonic and near-isometric muscle that identified the new pathway when probability for the 3 nm step-size was observed to exceeded probability for the 5 nm step-size. The latter is impossible for a 3 nm step coupled to a preceding 5 nm step (the 5+3 nm pathway) implying contribution from an independent solo-3 nm step-size pathway representing an additional complete transduction cycle *^28^*.

In the present work we use the Qdot assay to measure single βmys step-size and step-frequency characteristics when moving loaded cardiac and skeletal actin. Data classification constraints introduced with this application better separates signal from background giving higher signal-to-noise data from which we can resolve step-size probability densities in the sub-nanometer range. We find, for both skeletal and cardiac actin substrate, in vitro evidence for the solo-3 nm step-size pathway that is now shown to be a 4 nm step-size giving new direct evidence justifying its separation into a new pathway for myosin movement of actin. The solo-3^+^ nm step-size pathway (previously solo-3 nm step-size pathway) is occupied only under loaded conditions and is proposed to incorporate a clutch mechanism controlling myosin slip under load to facilitate the production of power under high resisting force. Our working model has cardiac myosin generating 8, 5, 5+3, and solo-3^+^ nm myosin step-sizes in the 4 pathway network where an autonomous myosin molecule regulates pathway choice at two points in the network with lever arm strain inhibited ADP release and a ratcheting ELC N-terminus strain sensor with selective resistance to movement in the direction of the loading force while permissive to movement in the direction of the contractile force. The lever arm mediates both strain sensing mechanisms.

Two actin isoforms, one dominant in skeletal the other in cardiac muscle, have a specific skeletal/cardiac actin stoichiometry of ~1:4 in a normal adult human heart *^29^*. It is likely that an increasing skeletal/cardiac actin stoichiometry is the programmed response of a disease-compromised heart to an inability to meet demand *^30^*. We demonstrate significant myosin step-size distribution differences for skeletal vs cardiac actin substrate because they differently tune βmys strain-dependent mechanics. We attribute cardiac actin impact on myosin load sensitivity to the unique and specific cardiac ELC N-terminus/cardiac-actin contact at Glu6/Ser358. It alters cardiac motor force-velocity characteristics detected with single myosin mechanics.

## METHODS

### Protein preparation

Bovine ventricular actin and rabbit skeletal actin were prepared and purified as previously described *^31^*. Briefly, myosin is extracted from minced muscle tissue with a high ionic strength neutral buffer solution containing ATP. The solubilized proteins including myosin are separated from the insoluble part of the muscle tissue including actin by centrifugation. Next, the actin-containing pellet is washed several times with acetone to denature and remove proteins other than actin. The remaining tissue fraction containing the actin is air-dried to form a powder. The actin powder is dissolved in a neutral low ionic strength buffer containing ATP. The dissolved actin is in the native monomer form called globular actin or G-actin. G-actin can be polymerized into its filamentous form (F-actin) by increasing the ionic strength. This property was used to further purify the extract by cycling between the G-and F-actin forms and using ultracentrifugation to separate the native monomer and polymer actin forms from the remaining impurities and denatured actin. The final preparation, at 99.9% pure as determined by sodium dodecyl sulfate polyacrylamide gel electrophoresis (SDS-PAGE, **Figure 1**), was stored immediately under argon gas in small aliquots in liquid nitrogen. The two actin isoforms migrate identically in the SDS-PAGE gel. Protein concentrations were determined by UV absorbance using A(1% @ 290 nm) = 6.4 for actin (42 kDa). Mass spectrometry on the tissue purified bovine cardiac ventriculum actin registers 22% skeletal and 78% cardiac actin *^32^* in excellent agreement with previous estimates using peptide sequencing of actin from the same tissue *^29^*. We refer to actin purified from ventriculum as cardiac actin.

**Figure 1.**
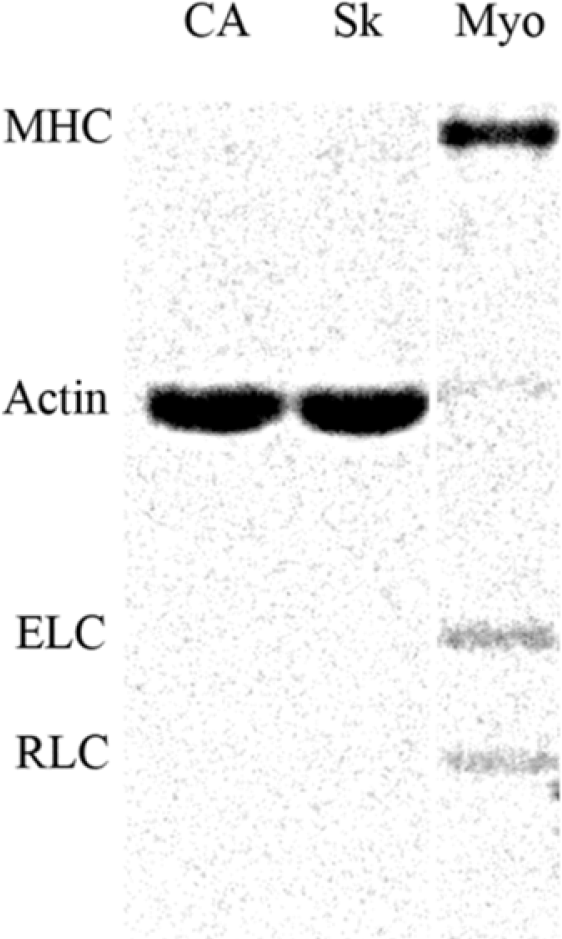
Figure 1. SDS-PAGE of cardiac (CA) actin, skeletal actin (Sk), and βmys (Myo). The column for βmys has regulatory and essential myosin light chains (RLC and ELC) bands indicated, and, contaminating actin. Baseline subtracted absorption integrated over the actin band is ~10% that of MHC.

Frozen G-actin was thawed and spun at 250,000×g for 90 min to remove denatured actin before use. Rhodamine labeling of actin filaments was performed with rhodamine-phalloidin and actin in a 1.2:1 molar ratio in a buffer containing 25 mM KCl, 0.2 mM CaCl_2_, 4 mM MgCl_2_, 10 mM DTT, 0.1 mM PMSF and 25 mM imidazole at pH 7.4 on ice for 12 h and then stored on ice until used within 5 days. Qdot labeling was performed as described previously *^33^*. Briefly, the actin filaments used for Qdot labeling were labeled with the mixture of biotin-xx phalloidin and rhodamine-phalloidin in a molar ratio of 1:9 and stored on ice. The overall molar ratio of phalloidin to actin and the buffer conditions were identical to that used for the rhodamine labeling.

Porcine βmys was prepared from porcine heart ventriculum as described^4, 33^. Briefly, βmys was extracted from minced, washed ventriculum for 10 min at 4 °C with “Guba-Straub” solution (0.3 M KCl, 5 mM ATP, 1 mM DTT, 5 mM MgCl_2_, 1 mM EGTA, 0.001 mg/ml leupeptin in 50 mM potassium phosphate buffer, pH 6.5). Solubilized myosin was separated from the tissue by centrifugation then three precipitation cycles eliminated contaminating soluble proteins. The pellet was then dissolved in a high salt buffer (0.6 M KCl, 5 mM ATP, 1 mM DTT, 5 mM MgCl_2_, 1 mM EGTA, 0.001 mg/mL leupeptin in 50 mM Tris-HCl pH 8.0 at 4 °C) followed by ultracentrifugation (250,000×g for 2 h). The upper 66% of the supernatant was collected and dialyzed overnight in storage buffer (0.6 M KCl, 2 mM DTT, 0.001 mg/mL leupeptin in 25 mM Tris-HCl pH 7.4 at 4 °C) followed by ultracentrifugation (250,000×g for 3 h) to remove remaining actin or actomyosin impurities. The βmys was stored in sealed tubes at −20°C in 50% glycerol (vol/vol). Purified βmys has ~10% actin impurity judged from analyzing the SYPRO Ruby stained SDS/PAGE gel shown in **Figure 1**. Urea gel electrophoresis and mass spectrometry analysis of the βmys did not detect phosphorylated RLC in the βmys used in experiments *^34^*.

### Actin-activated myosin ATPase

βmys stored in 50% glycerol was precipitated with addition of 12 volumes of ice-cold water containing 2 mM DTT, collected by centrifugation, and then resuspended in 300 mM KCl, 25 mM imidazole (pH 7.4), 4 mM MgCl_2_, 1 mM EGTA, 10 mM DTT, and 0.01 mg/mL leupeptin. Myosin at a final concentration of 0.6 μM was titrated with 1, 3, 5, 9, 22, 50, and 93 μM F-actin. The ATPase assay buffer contained 25 mM imidazole (pH 7.4), 4 mM MgCl_2_, 1 mM EGTA, 10 mM DTT, 0.01 mg/mL leupeptin, and a final KCl concentration of 25 mM. ATPase reaction was initiated by addition of 3 mM ATP, and the mixture was incubated at 21 °C for 5 min. Inorganic phosphate measurements were performed using the Fiske and Subbarow method *^35^*.

Actin-activated ATPase results were characterized using Michaelis-Menten kinetics,

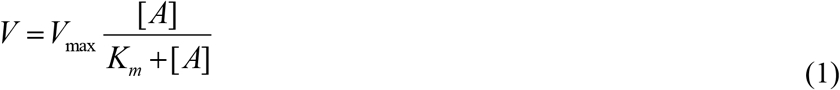

for actin concentration [A], V the ATPase rate equal to V_max_ at saturating actin, and actin binding constant K_m_. V vs [A] curves were fitted using a nonlinear algorithm to determine constants V_max_ and K_m_.

### In vitro motility and Qdot assays

In vitro motility and Qdot assays were performed in a flow cell using total internal reflection fluorescence (TIRF) microscopy exactly as described *^23^*. Motility buffer included 25 mM KCl, 25 mM imidazole (pH 7.4), 4 mM MgCl_2_, 1 mM EGTA, 20 mM DTT, 0.01 mg/mL leupeptin, 0.7% methylcellulose, 2 mM ATP, 3 mg/mL glucose, 0.018 mg/mL catalase, and 0.1 mg/mL glucose oxidase. The flow cell was infused at the start with 0.04-0.4 μM myosin. Actin sliding velocities for the in vitro motility assay, s_m_, and the length of actin filaments were quantitated using FIESTA software *^36^*.

Frictional loading assays were performed like the unloaded assays except that the flow cell was infused at the start with the mixture of myosin and α-actinin in concentrations of 0.16 μM myosin and 0-7 μg/mL α-actinin (Cytoskeleton, Denver, CO).

In the Qdot assay, images were acquired with an EMCCD camera (Andor, Belfast, UK) in 45 ms intervals indicated by Δt and using Andor’s SOLIS software. Each movie recorded images for 36 seconds. Intensity values were converted to photons using the conversion formula in SOLIS and the images output in TIFF format for reading into ImageJ. We tracked movement of the Qdot labeled actin at super resolution using the ImageJ plugin QuickPALM *^37^*. Baseline histograms corresponding to thermal/mechanical fluctuations were recorded likewise from Qdot labeled actin immobilized on the surface by myosin in the absence of ATP. Thermal/mechanical fluctuation baseline contributions to active myosin event-velocity histograms were estimated as described below under *Qdot Assay Event-Velocity Histogram Simulation*. All *in vitro* motility and Qdot assay experiments were conducted at room temperature (20-22 °C).

### Force calibration in the loaded actin in vitro motility and Qdot assays

Loaded *in vitro* motility and Qdot assay data were fitted to a viscoelastic model of frictional loads developed by Greenberg & Moore *^38^* and as described previously *^25^*. Average sliding filament motility velocity, s_m_,

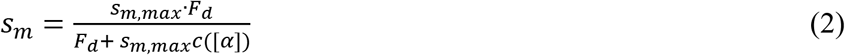

has s_m,max_ the velocity at zero load, F_d_ the ensemble driving force, and friction coefficient c([α]),

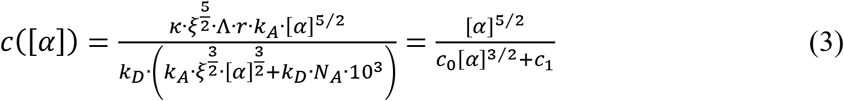

for κ system compliance associated with α-actinin (1.7 pN/nm), ξ a constant defining the surface geometry of α-actinin on the flow cell surface (3.97×10^21^ M^−1^ m^−2^), r the reach of α-actinin to bind to the actin filament (82 nm), k_A_ the second-order rate constant for α-actinin attachment to the actin filament (4×10^6^M^−1^s ^−1^), [α] molar concentration of α-actinin, and k_D_ α-actinin detachment rate (9.6 s^−1^). The constants are suggestions based on the literature and identical to those already described *^38^*. Other known quantities in eq. 3 are Avogadro’s number, N_A_, and actin filament length in our system, Λ ≈ 1 μm. Coefficients c_0_ and c_1_ in eq. 3 are optimized when fitting data (discussed below) thus providing an experimental value for their relationships to the constants quoted above and implied by eq. 3.

Frictional force, F_f_, exerted by the α-actinin in the loaded motility assay,

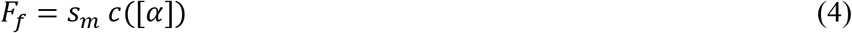

relates force and velocity in the Hill equation *^39^* for normalized force and velocity,

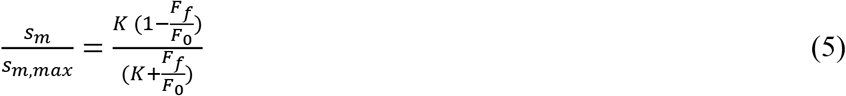

and implies power output, P,

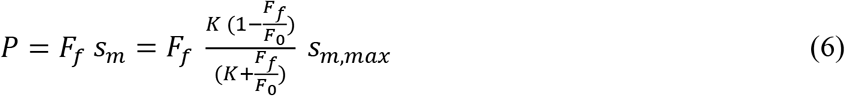

for F_0_ isometric force and dimensionless constant K proportional to the myosin attachment rate f_APP_ *^40^*. Assuming F_d_ = F_0_, we estimate free parameters F_0_, *K*, c_0_, and c_1_ using linear programming with constraints from the raw data consisting of [α] vs s_m_ data points for skeletal and cardiac actin, eqs. 2–5, and requiring all fitted parameters are ≥ 0. Fitted constants F_0_ and K have independent values for skeletal or cardiac actin while c_0_ and c_1_ are identical for both species. In addition, the maximum unitary force generated by βmys is likely ~2 pN *^41^* and the number of βmys simultaneously impelling a 1μm long (skeletal) F-actin is ~15 *^34^* implying an upper limit constraint on F_0_ of 60 pN (twice the estimated maximum of 30 pN). When solved, parameters calibrate the frictional force (in pN) using eqs. 3 & 4 and give best estimate fitted curves for s_m_ vs [α] (eqs. 2 & 3), s_m_ vs F_f_ (eq. 5), and P vs F_f_ (eq. 6).

### Qdot Assay Event-Velocity Histogram Simulation

*In vitro* motility has the myosin moving actin with a motility velocity s_m_ such that,

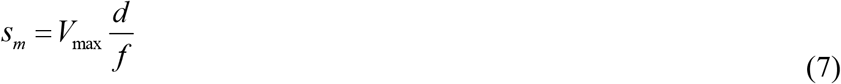

for myosin unitary step-size *d* and *f* the duty ratio given by the time actomyosin is strongly bound during an ATPase cycle, t_on_, divided by the cycle time, 1/V_max_, hence,

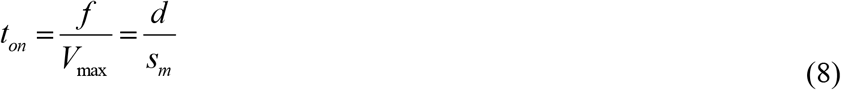

Myosin executes one of three unitary steps with step-size, d_j_, and relative step-frequency, ω_j_, for unitary step *j* = *S, I*, and *L* where S, I, and L are for the short (~3-4 nm depending on load), intermediate (~5-6 nm depending on load), and long (~8 nm) nominal unitary steps of cardiac myosin *^33, 42^*.

Relative step-frequency is a characteristic proportional to the rate of cross-bridge cycling with the higher rate producing a more frequent *j* ^th^ step with step-size d_j_. The dimensionless relative step-frequency is normalized with ω_S_+ω_I_+ω_L_=1. The absolute cycling rate for step *j*, V_j_, has V_j_ = V_max_ ω_j_ and with *V*_*max*_ = Σ_*j*=*S,I,L*_ *V*_*j*_.

In an ensemble of cross-bridges interacting with one actin filament, like the conditions in every muscle or motility assay, only one actin velocity is possible hence motility velocity s_m_ is the same for each unitary step-size implying each step-size has unique duty ratio and time strongly actin bound. From eq. 8, step *j* duty ratio,

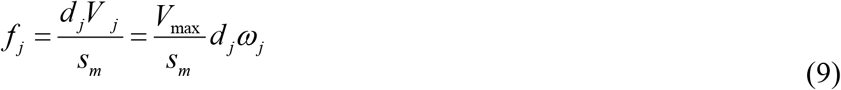

From eqs. 8 & 9, the time myosin spends strongly bound to actin,

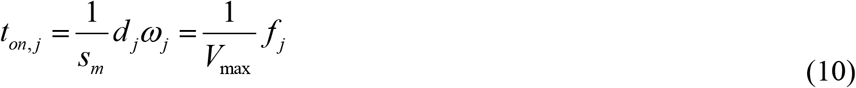

Eq. 10 shows that each unitary step has a unique t_on_ that varies with step-size and relative step-frequency. The t_on_ is distributed exponentially as observed using the laser trap for the in vitro actin detachment rate *^43^*. The ensemble average quantities used include average step-size,

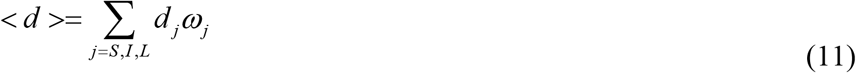

and average duty-ratio <f>,

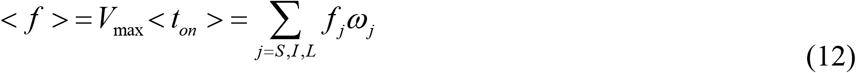

Simultaneously bound cross-bridges produce identical actin sliding velocity, s_m_, for step-size, d, and the time the cross-bridge is actin bound, t_on_, such that s_m_ = d/t_on_. Net velocity is proportional to the time any cross-bridge is actin bound between frames. At higher velocities, when multiple cross-bridges impel actin, it is more likely that two or more will overlap some of their bound time, producing net sliding velocity intermediate to the discrete velocities even within the domain of the unitary step-size velocity. This effect is seen as the rising baseline in the panels that becomes more significant at higher resisting force with rising <t_on_> due to strain.

We simulated motility assay event-velocity histograms using a time × space array representing an actin filament interacting with the surface-bound myosin. Array rows represent myosin binding sites on actin located every 36 nm along the filament that interact with surface bound myosins while columns represent their time evolution. We simulated a ~1 μm long actin filament having 28 total myosin binding sites. The time × space array was filled one row at a time by randomly generating binding site occupation from actin binding probability *p*.

We partitioned t_on_ into *n* segments (t_seg_ = t_on_/n). An occupied binding site remains occupied for several t_seg_’s (one t_seg_ elapses between rows in the time × space array) and on average for <t_on_>. The total array column length is equal to Δt/t_seg_, for Δt, the time interval between consecutive frames. The actin filament velocity is then the number of rows with at least one site occupied times the myosin step size divided by nΔt. The simulation for a single actin filament and for Δt = 45 ms is run repeatedly until the number of events falling within the unitary step-size domain approximately matches the number observed.

Experimental data, **v**_obs_ and **v**_t/m_, and simulated data, **v**_sim_, are event-velocity histograms in a vector representation. The **v**_obs_ is the observed velocities of Qdot labeled actin in the presence ATP. The **v**_t./m_ velocities quantitate actin movement from thermal/mechanical fluctuations measured independently from Qdot labeled actin immobilized on the surface by myosin in the absence of ATP. The **v**_sim_ simulated velocities quantitate the actin movement from active myosins in the model just described. These event-velocity histograms are linearly related by,

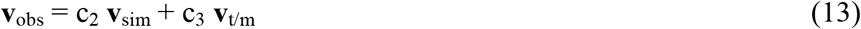

for **v**_sim_ and **v**_t/m_ **ℓ**_*1*_ normalized vectors, and, c_2_ and c_3_ unknown scalar constants equal to the total myosin-based and thermal/mechanical-based actin displacement events. Unknowns c_2_ and c_3_ are solved optimally using linear programming under the equality constraint that known total observed events = c_2_ + c_3_ and the inequality constraints c_2_ ≥ 0 and c_3_ ≥ 0.

V_max_ and motility velocity, s_m_, are measured under saturating actin and myosin conditions, respectively. They are constant parameter inputs to the simulation that are characteristic to each actin tested. Simulation approximates the Qdot motility event-velocity histogram in the low velocity domain of 0~4 natural velocity units (vu) where 1 vu = (d_I_/Δt) for dI the intermediate step-size (near 5 nm) and where unitary events dominate. The unknown parameter set actively searched in the simulation consists of the actin binding probability for myosin, *p* (1 free parameter), step-size (3 free parameters), and relative step-frequency (3-1 = 2 free parameters due to normalization). Trial parameter values are generated in the simulation by random choice from a range of values set at the start of the simulation as described below. Simulations sampled all of parameter space.

Peak position and area for the first three peaks in event-velocity histograms are the unitary step-size and relative step-frequency estimates obtained directly from inspection of the data. We observed 3 normally distributed step-sizes with standard deviations (SDs) within 0.3-0.7 nm, and, 3 normally distributed step-frequencies and SDs within 0.1-0.3. We searched step-size and step-frequency parameter space in 5 dimensions over a range defined by ±1.5 standard deviations in 7 intervals. The actin binding probability for myosin and its parameter space dimensions were estimated empirically and spanned in 7 intervals during the search for best fitting simulations. Estimates for step-size and step-frequency from simulation rigorously account for step overlap when two or more myosins impinge on one actin filament during some of the time they are strongly actin bound.

**Figure 2.**
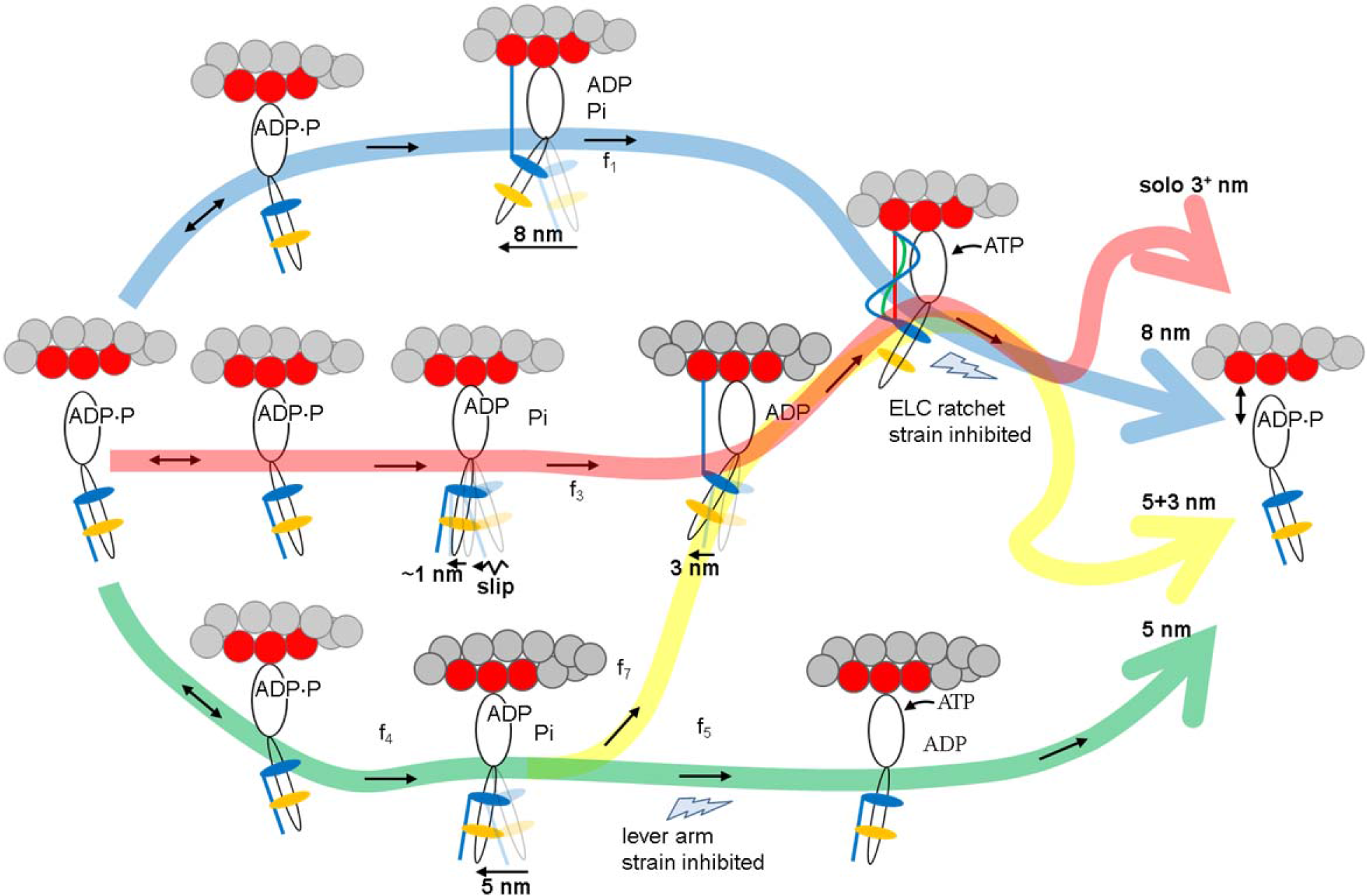
The 4 pathway network for cardiac myosin unitary steps. An unloaded myosin powerstroke generally has two sequential enzymatic steps with Pi release driving the larger lever arm swing (nominal 5 nm step-size) followed by the ELC N-terminus binding actin and ADP release driving the smaller lever-arm swing (3 nm ste-size). The blue pathway has an 8 nm step-size where large and small steps are tightly coupled for the maximum lever-arm swing. The green pathway is predominant with the 5 nm step accompanying product release. Occasionally Pi release with the 5 nm step is not immediately followed by ADP release because of lever arm strain inhibition at the lower thunderbolt allowing time for the subsequent ELC N-terminus binding and 3 nm step in the yellow pathway. The latter is the least likely pathway for unloaded myosin. Flux through the network pathways is variously distorted by loading. The red pathway denotes a solo-3^+^ nm step-size where myosin partially slips to releases Pi with a ~1 nm net forward movement but then the ELC N-terminus binds actin permitting ADP release and completion of a 3 nm step. Myosin in the red pathway is observed here as a new 4 nm step-size occurring only in a high-force regime defined in the text. The ELC-ratchet strain inhibiting filter at the upper thunderbolt regulates the 3 nm part of the 5+3 nm pathway as well as the solo-3^+^, and 8 nm pathways (yellow, red, and blue). ELC ratchet strain inhibits ATP binding and ELC N-terminus detachment from actin maintaining tension at peak isometric force. The lever arm strain inhibition at the lower thunderbolt regulates the 5 nm pathway (green) with strain inhibition of ADP release. The upper and lower thunderbolts are competing strain regulated checkpoints that modulate step-frequencies for quickly responding to changing force-velocity demands. Myosin flux through the network is denoted in the figure with f_i_ for i = 1, 3, 4, 5, and 7, and identically to earlier work *^25^*. Flux calculations were not attempted here because the constraints limiting the flux values are sometimes qualitative as described in the text.

### 4-pathway contraction model

Earlier work using the unloaded in vitro Qdot assay on βmys *^33^* combined with the in vivo single cardiac myosin imaging of zebrafish embryo hearts *^28^* suggested a 4 pathway network summarized in **Figure 2** for generating 8 (blue pathway), 5 (green), 5+3 (green-yellow), and solo-3^+^ (red) nm myosin step-sizes *^25^*. In **Figure 2**, weak actin binding myosin reversibility (indicated with ↔) preemptively avoids pathways that do not complete cycles under increasing load. Under loaded conditions, flux through the cycle is checked at two strain inhibited points. The traditional lever arm strain checkpoint inhibits ADP release and hence ATP dissociation of actomyosin following the 5 nm step-size (green path) sometimes sending flux towards the 3 nm step-size (yellow branch). The ELC-ratchet strain checkpoint regulates the solo-3^+^, 8, and 5+3 nm pathways (red, blue, and green-yellow). The latter senses tension in the actin bound ELC N-terminus such that the slack ELC extension (blue, when muscle is unloaded or rapidly shortening under low load) allows ATP binding and quick detachment from actin to complete the cycle, the moderately tense ELC extension (green, when muscle is under moderate loads approaching isometric) partially inhibits ATP binding and detaches slower from actin to exert static myosin tension contributing to force on slowly translating muscle filaments, and the maximally tense ELC extension (red, when muscle is near isometric) inhibits ATP binding and detaches slowest from actin to exert static myosin tension contributing more of the force on the static muscle filaments. The ELC-ratchet tends to filter out longer step-sizes as tension rises since they will transfer a larger tension to it. Both the traditional lever arm strain checkpoint inhibiting ADP release, and the ELC-ratchet strain checkpoint regulation, are mediated by lever arm strain.

Strongly bound myosins contributing static tension are the force bearing 0 length step-size myosins not explicitly depicted in **Figure 2** [see rather **Figure 8** in *^28^*]. Modulating flux through 2 strain-dependent steps with different inhibitions adjusts average step-size. The favored pathway at high tension (>80% of cycles) involves the solo-3^+^ nm step via the red pathway. It releases Pi with slight net forward movement probably by slipping at high tension but then completes a 3 nm step. Results introduced here suggest slippage of 4 nm extends the solo-3^+^ nm step-size to 4 nm. Slip distances of 2-8 nm were observed in synthetic myofilaments under tension (see Fig 1 panels e-g in *^44^*, larger slips were observed earlier *^45^*). This pathway is populated even at the highest tension implying the ELC ratchet strain checkpoint is less inhibiting than the ADP release inhibiting checkpoint when in near-isometric contraction *^46^*.

### Homology modelling

A molecular model for the skeletal myosin S1(A1) isoform bound to skeletal F-actin was described with 14 actin monomers forming the actin filament and two S1’s with light chains bound to the filament *^18^*. The model is based on crystal structures for skeletal myosin (2mys) *^47^* and skeletal actin (1atn) *^48^*, and, electron density maps for S1 bound to F-actin *^49–51^*. The ELC(A1) N-terminal extension is missing from the model structures and was constructed using structural pattern searches and energy minimization. Three actin monomers contacting S1 (two contacting the heavy chain and another contacting the ELC N-terminus) were extracted from that structure to use as the model for the all-cardiac isoform structure. Cardiac actomyosin was homology modeled using Modeller *^52^* and human sequences ACTC1, MYH7, MYL2 (RLC), and MYL3 (ELC).

### Statistics

Actin-activated ATPases were measured 4 times with at least 2 different actin and 2 different βmys preparations. Significance testing generally utilizes one-way or two-way ANOVA with Bonferroni or Tukey-Kramer post-tests for significance.

Actin motility velocity data sets consist of 7-31 acquisitions (one acquisition is one *in vitro* motility movie) per actin isoform and α-actinin concentration. Qdot assay data sets consist of 8-16 acquisitions per actin isoform and α-actinin concentration. Data sets for motility velocity and Qdot assays were accumulated from at least 2 different actin and 2 different βmys preparations.

We simulated data for each event-velocity histogram independently to estimate single myosin mechanical characteristics consisting of 3 step-sizes and 3 step-frequencies. We compared fitted parameters as categorical variables in factor 1 for the 8-16 independent event-velocity histograms in a data set in factor 2 using two-way ANOVA with Bonferroni or Tukey-Kramer post-tests for significance at the 0.05 level. This test indicated no significant difference among event-velocity histograms in data sets hence parallel acquisitions were pooled.

Simulated data ensembles were created by using best fitting event-velocity histogram simulations generated for a Qdot assay data set where simulations are combined linearly to approximate the event-velocity histogram from the pooled data with coefficients ≥0 while minimizing the χ^2^ goodness-of-fit test with all points equally weighted. The number of best fitting simulations combined is ≤ the number of independent event-velocity histograms in a data set.

### Effect of actin mixtures

Actin purified from rabbit skeletal muscle is the homogeneous skeletal isoform *^53^* while that from bovine ventriculum contains 22 and 78% skeletal and cardiac isoforms, respectively *^32^*. Measured characteristics using actin-activated ATPase, actin motility, and Qdot labeled actin motility contrast responses of pure skeletal and heterogeneous cardiac actin samples. We show that, for the two actin species, actin-activated ATPase is statistically identical while other quantities are significantly different.

We investigated the effect of actin mixtures on motility velocity by testing sensitivity of parameters estimated from the motility using actin velocity vs loading force data corrected by,

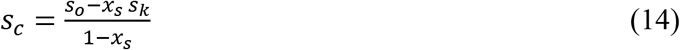

where x_s_ is the fraction of skeletal actin in the mixture (~0.22), s_c_ and s_k_ pure unknown cardiac and known skeletal actin isomer velocities vs loading force, and s_o_ the observed actin velocity vs loading force from ventriculum purified actin. We similarly estimated the effect of the actin mixture on Qdot assay measured averaged quantities including filament velocity, step-size (<d>), and average duty-ratio (<f>) using eq. 14 and substituting *s* with the mentioned Qdot assay measured quantity.

Presumption of eq. 14 has actin monomer subunits impacting velocity independently when the isoforms comingle in cardiac F-actin. We show below that it is a reasonable supposition for this system.

## RESULTS

### Actin-activated ATPase of cardiac myosin

**Figure 3** shows the actin-activated myosin ATPase vs actin concentration, [A], for skeletal (red triangles) and cardiac actin (black circles) with error bars indicating standard deviation. Curves were fitted using eq. 1 to obtain the Michaelis-Menten parameters V_max_ = 0.942 ± 0.104 s^−1^, K_m_ = 6.8 ± 2.7 μM for cardiac actin, and, V_max_ = 0.804 ± 0.106 s^−1^, K_m_ = 4.7 ± 2.5 μM for skeletal actin and as indicated in **Table 1**.

**Figure 3.**
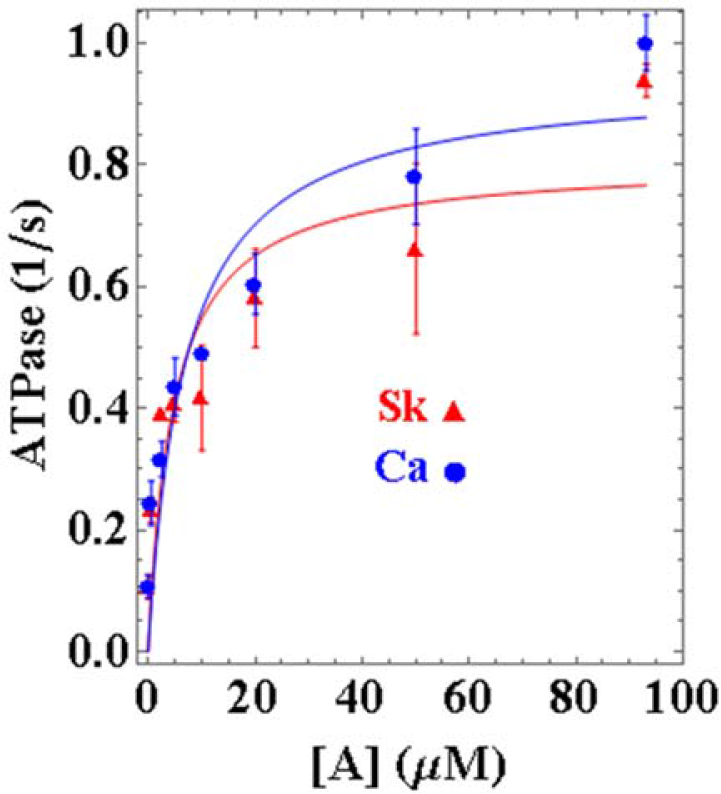
Actin-activated βmys ATPase in inverse seconds vs actin concentration [A] in μM for cardiac (Ca blue) and skeletal (Sk red) actin. Error bars show standard deviation for sampling statistics given in Table 1 and under experimental conditions given in Methods. Fitted curves use eq. 1. No significant difference is detected between the ATPase rate vs [A] for cardiac and skeletal actin.

**Table 1:** Mechanical characteristics of βmys for two actins

| **Table 1: Mechanical characteristics of βmys for two actins <sup>1</sup>** |  |  |  |
| --- | --- | --- | --- |
|  | Skeletal actin | Cardiac actin | p< |
|  | Actin-activated myosin ATPase |  |  |
| $V_{\max}$ (1/s) | 0.80±0.10 (N=4) | 0.94±0.10 (4) | ns @ 0.05 |
| $K_m$ (μM) | 4.7±2.5 | 6.8±2.7 | ns @ 0.05 |
|  | Peak in vitro motility velocity for unloaded actin |  |  |
| $S_{m,\max}$ (μm/s) | 0.20±0.01 (N=31) | 0.16±0.01 (23) | 0.01 |
| | Isometric force ($F_0$) in pN & dimensionless attachment rate (K) for motility data | | |
| $F_0$ | 18.4±3.7(N=12) | 14.3±2.8 (12) | 0.01 |
| K | 10.1±4.2 | 7.0±4.0 | 0.01 |
| $F_0$ corrected | 21.0±2.3 (12) | 15.3±1.5 (12) | 0.01 |
| K corrected | 10.3±4.0 | 3.7±2.5 | 0.01 |
| | Isometric force ($F_0$) in pN & dimensionless attachment rate (K) for Qdot assay motility data | | |
| $F_0$ | 38.5±9.8 (N=8) | 29.5±10.5 (8) | 0.01 |
| K | 18.0±0.1 | 14.0±5.1 | 0.01 |
| $F_0$ corrected | 36.5±9.4 (8) | 29.1±12.3 (8) | 0.01 |
| K corrected | 18.0±0.1 | 12.0±6.1 | 0.01 |

<sup>1</sup> Average values and standard deviation indicated for (N) replicates. ANOVA significance under the p< heading indicates confidence level for distinguishing skeletal from cardiac actin or ns for not a significant difference. $F_0$ or K corrected refers to optimization of eqs. 2–6 using corrected cardiac actin data from eq. 14.

We tested significance of data in **Figure 3** using 2-way ANOVA with factor 1 proteins skeletal actin and cardiac actin, and, factor 2 the actin concentration [A]. Differences between actin-activated myosin ATPases for the two actin isoforms are not significant for confidence level p < 0.05.

Cryo-EM data indicates rigor actomyosin contacts (in the presence of tropomyosin) between myosin Loop 2 and N-terminal actin residues 1-4 *^54^*. The actin N-terminus contains skeletal/cardiac actin substitutions Glu2Asp and Asp3Glu while Loop 2 links the 50 and 20 kDa proteolytic fragments of S1. Loop 2 substitutions dramatically impact actin-activated ATPase and actin binding affinity *^55, 56^* implying Loop2/actin N-terminus contacts participate integrally in normal motor function, and, the possibility for skeletal vs cardiac actin motor function differences. However, data in **Figure 3** implies the Loop2/actin interaction is negligibly impacted by the skeletal/cardiac actin Glu2Asp and Asp3Glu substitutions.

A single cardiac myosin contacts 3 actins while the ELC N-terminus extension binds just one. Assuming Glu2Asp and Asp3Glu substitutions do not affect the actomyosin interaction, each βmys has either a cardiac or skeletal actin-type contact but never a hybrid of forms. Then the average number skeletal actomyosin contacts during motility are linear with the amount of skeletal actin and justifying the presumption of eq. 14.

### Loaded motility

**Figure 4 panel a** shows βmys in vitro motility velocity, s_m_, vs α-actinin concentration, [α], for cardiac (Ca, blue circles) and skeletal actin (Sk, red triangles). **Panel b** shows the same data in velocity vs normalized frictional force described by the Hill equation (eq. 5). **Panel c** indicates the same data in a power vs normalized frictional force representation using eq. 6. βmys generates ~2 fold more peak power when moving skeletal compared to cardiac actin.

**Figure 4.**
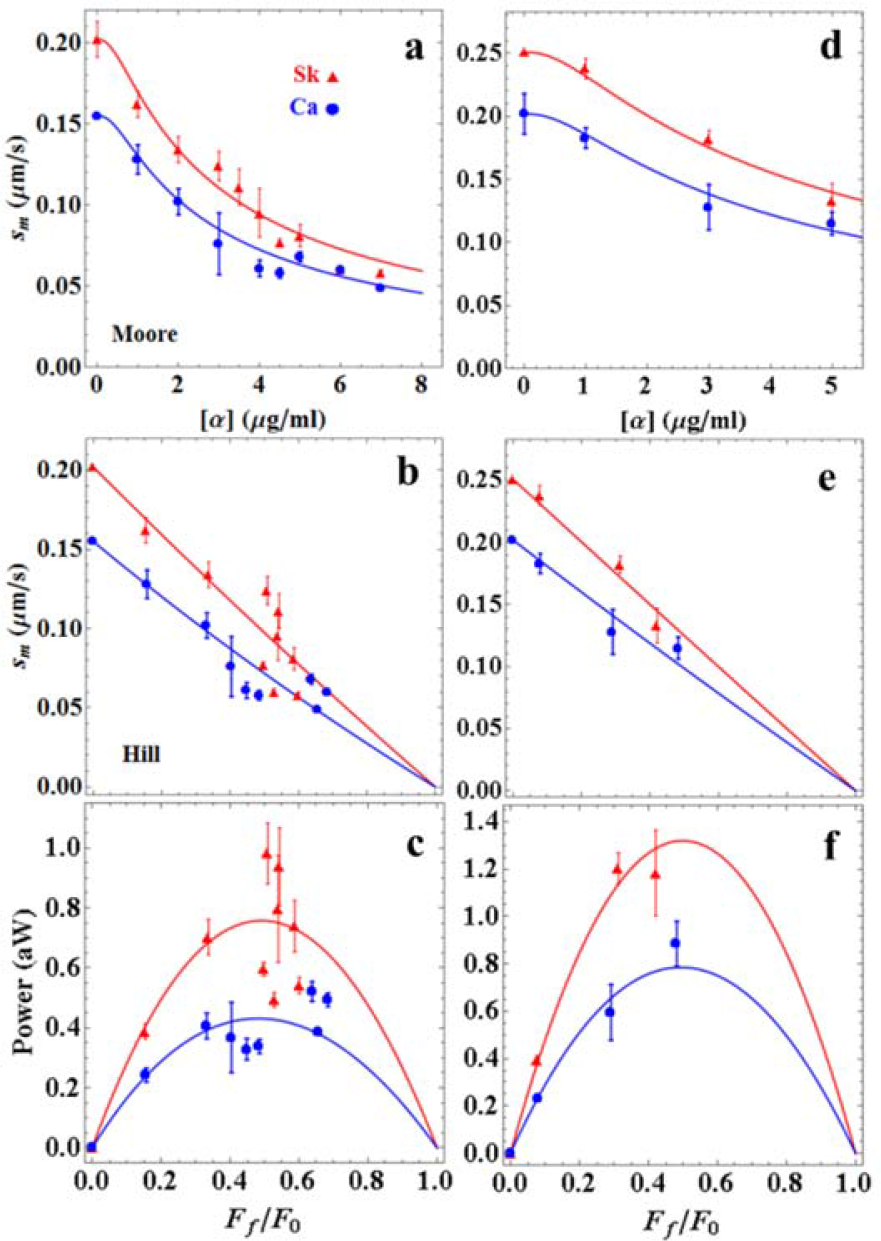
βmys loaded in vitro motility assay for skeletal (Sk red) or cardiac (Ca blue) actin. Panel a: Motility velocity, s_m_, vs α-actinin concentration. Panel b: Motility velocity as in panel a vs normalized frictional force F_f_/F_0_ for F_f_ the frictional loading force and F_0_ isometric force. Panel c: power vs F_f_/F_0_. Error bars show standard deviation for 7-31 acquisitions at each α-actinin concentration and under experimental conditions given in Methods. Fitted curves are based on eqs. 2–6 applied to data as described in Methods. Experimental data appears as discrete points with error bars and fitted curves as solid lines. Significance testing of cardiac vs skeletal data sets in panels a–c indicates they differ significantly in each case with confidence level p < 0.01. Panels d–f are identical to panels a c except for Qdot labeled actin under fewer and slightly different loads and with error bars showing standard deviation for 8-16 acquisitions.

Error bars indicate standard deviation for 7-31 acquisitions at each α-actinin concentration. We tested significance of s_m_ vs [α] data (**panel a**), s_m_ vs F_f_/F_0_ data (**panel b**), and power vs F_f_/F_0_ data (**panel c**) using 2-way ANOVA with factor 1 the skeletal or cardiac actin, and, factor 2 the [α] or F_f_/F_0_ values. Data differs significantly with confidence level p < 0.01.

Skeletal and cardiac actin fitted curves in **Figure 4 panels a-c** are optimized with free parameters F_0k_, F_0c_, K_k_, K_c_, c_0_, and c_1_ constrained in eqs. 2–6 where F_0k_ and F_0c_ are isometric forces for myosin moving skeletal and cardiac actin, respectively. Dimensionless constants K_k_ and K_c_ for skeletal and cardiac actin (eq. 5) are proportional to myosin attachment rate f_APP_. We find skeletal actin supports a significantly larger isometric force than actin purified from cardiac ventriculum. K_k_ and K_c_ are 10.1 and 7.0 for the skeletal and cardiac actin and are ~40x larger than that reported from intact slow skeletal muscle *^40, 57^*. Rate f_APP_ distinguishes the actin isoforms suggesting the skeletal actin promotes the force generating state as also implied by power in **panel c**. Mean and standard deviation values for F_0k_, F_0c_, K_k_, and K_c_ are summarized in **Table 1**. Fitted parameters c_0_, and c_1_ reflect on the various quantities defining the friction coefficient in eq. 3. They suggest that the values mentioned in Methods contributing the ratios, k_D_ k_A_ ξ^3/2^ and k_D_^2^ N_A_ 10^3^ over κ ξ^5/2^ Λ r k_A_, and corresponding to constants c_0_ and c_1_ in eq. 3 underestimates their true value in the motility assay by ~100 fold.

We estimated the impact of the skeletal actin component in ventriculum purified actin using corrected cardiac actin velocity data from eq. 14, constants c_0_ and c_1_ from the fitted curves in **Figure 4 panels a-c** (i.e., the implicit assumption that k_D_, k_A_, ξ, κ, r and Λ are equal for skeletal and cardiac actin) and with free parameters F_0k_, F_0c_, K_k_, and K_c_ optimized while constrained by eqs. 2–6. We find the free parameters do not differ significantly for cases using observed or corrected cardiac actin velocity data. Results for corrected cardiac actin velocity data are also indicated in **Table 1**. The skeletal actin impurity in the actin purified from ventriculum has negligible impact on in vitro motility.

Qdot labeled skeletal and cardiac actin velocity data in **Figure 4 panels d–f** parallel those in **panels a c** except that [α] sampling is limited to 0, 1, 3, and 5 μg/ml. Skeletal and cardiac actin fitted curves in **Figure 4 panels d–f** were surmised by methods identical to those used to fit the data in **Figure 4 panels a–c**. We find the Qdot labeling alters slightly the α-actinin/actin interaction reflected in modest changes in the optimized values for c_0_, and c_1_ while other fitted parameters are practically identical but with larger standard deviation estimates. Mean and standard deviation values for F_0k_, F_0c_, K_k_, and K_c_ are summarized in **Table 1**.

The Qdot labeled actin data was collected for the single myosin experiments, however, in their alternate form summarized by **Figure 4 panels d f** we use it to evaluate the impact of the skeletal actin component in ventriculum purified actin on Qdot labeled actin motility. Results compared in **Table 1** show that the fitted parameters pertaining to myosin function do not differ significantly for the observed or corrected (using eq. 14) cardiac actin velocity data. This ensemble measurement shows that isometric force and actin attachment rate in eqs. 2–6 for ventriculum purified actin are negligibly impacted by the small skeletal actin component suggesting pooled single myosin characteristics measured with the Qdot assay are also likely to be negligibly impacted.

### Loaded Qdot assay

The Qdot assay has Qdot labeled actin filaments ~1 μm long translating over surface bound βmys. **Figure 5** shows the Qdot assay pooled data event-velocity histogram from 8-16 acquisitions (N) for βmys moving actin from cardiac (left column) or skeletal (right column) muscle and for increasing frictional load indicated in pN. Event-velocity histogram domains cover 0~4 natural velocity units (vu) where (d_I_/Δt) = 1 for d_I_ the intermediate step-size, usually ~5-6 nm, and frame capture interval Δt = 45 ms. Summary data (solid squares connected by dashed line) has the baseline contribution from thermal/mechanical fluctuations already subtracted to show motility due to myosin activity. Peaks or inflection points below 2 vu are short (short red up arrow or S), intermediate (d_I_ and longer green down arrow or I), and long (longest blue up arrow or L) unitary step-sizes in nm. Green and blue arrows also indicate where some of the unitary step combinations should fall. Simulation of each event-velocity histogram acquisition is summarized for the pooled data by the solid line overlaying summary data.

**Figure 5.**
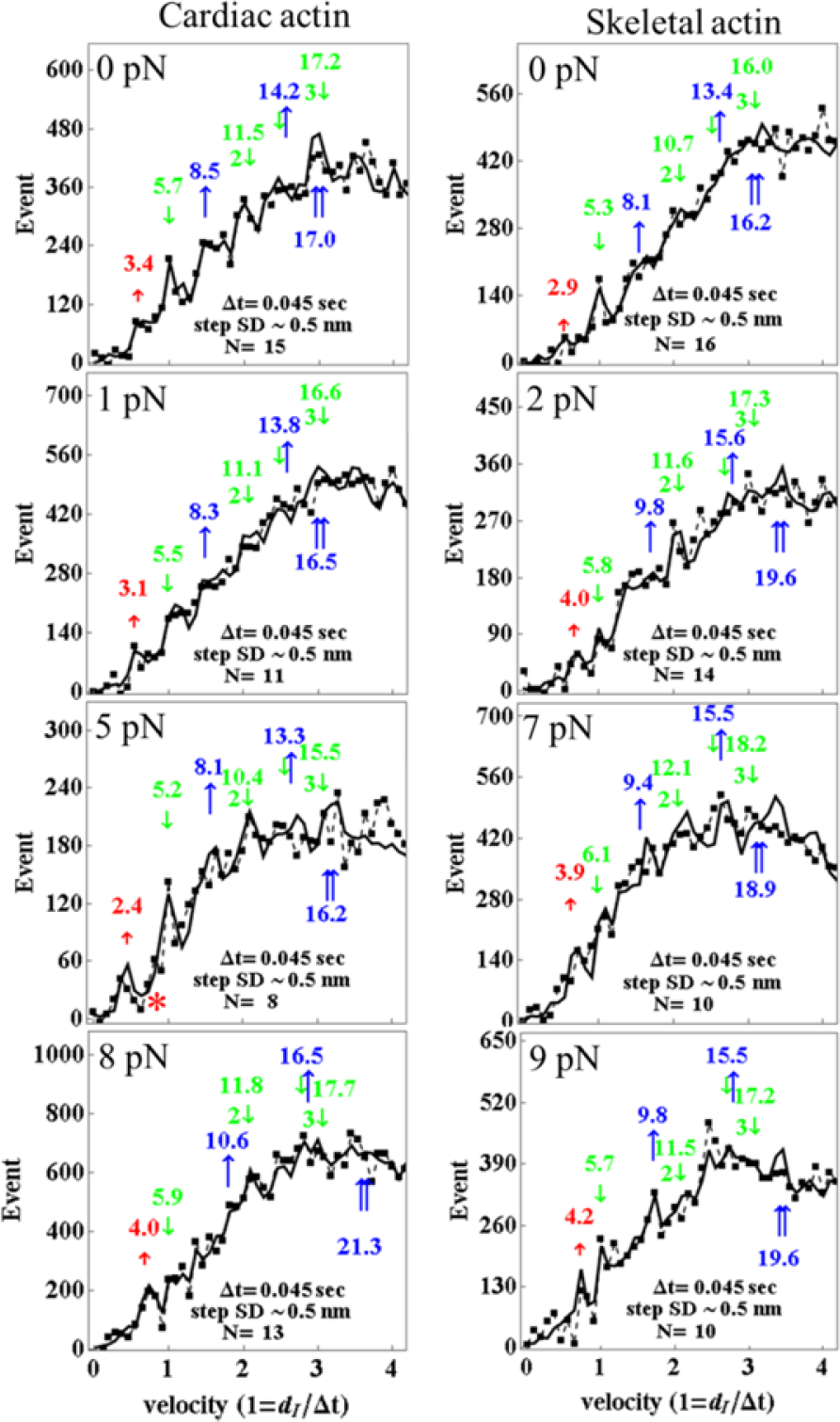
Event vs velocity histograms (solid squares) for βmys moving cardiac (left column) or skeletal (right column) actin and for frictional actin loading, F_f_, from 0 to 9 pN. Black solid lines are simulations performed as described in Methods and used to estimate step-size (at arrows) and step-frequency in Figure 6. Natural velocity units (vu) have 1 vu = (d_I_/Δt) for d_I_ the intermediate step-size (green down arrow at ~ 5-6 nm) between the short (red up arrow at ~ 2-4 nm) and long (blue up arrow at ~ 8-9 nm) step-sizes. Step-sizes have standard deviation of ~0.5 nm for the 8-16 replicates.

**Figure 6** shows the step-frequency expectations (lines), expectation values and their standard deviations for N acquisitions for the S, I, and L unitary steps derived from simulation of each event-velocity histogram acquisition. The expectation curves indicate the relative probability for step-frequency along the abscissa. The area under the color coded curves for the S, I, and L steps equals expectation values ω_S_, ω_I_, and ω_L_, respectively. The sum ω_S_ + ω_I_ + ω_L_ = 1 in each panel.

**Figure 6.**
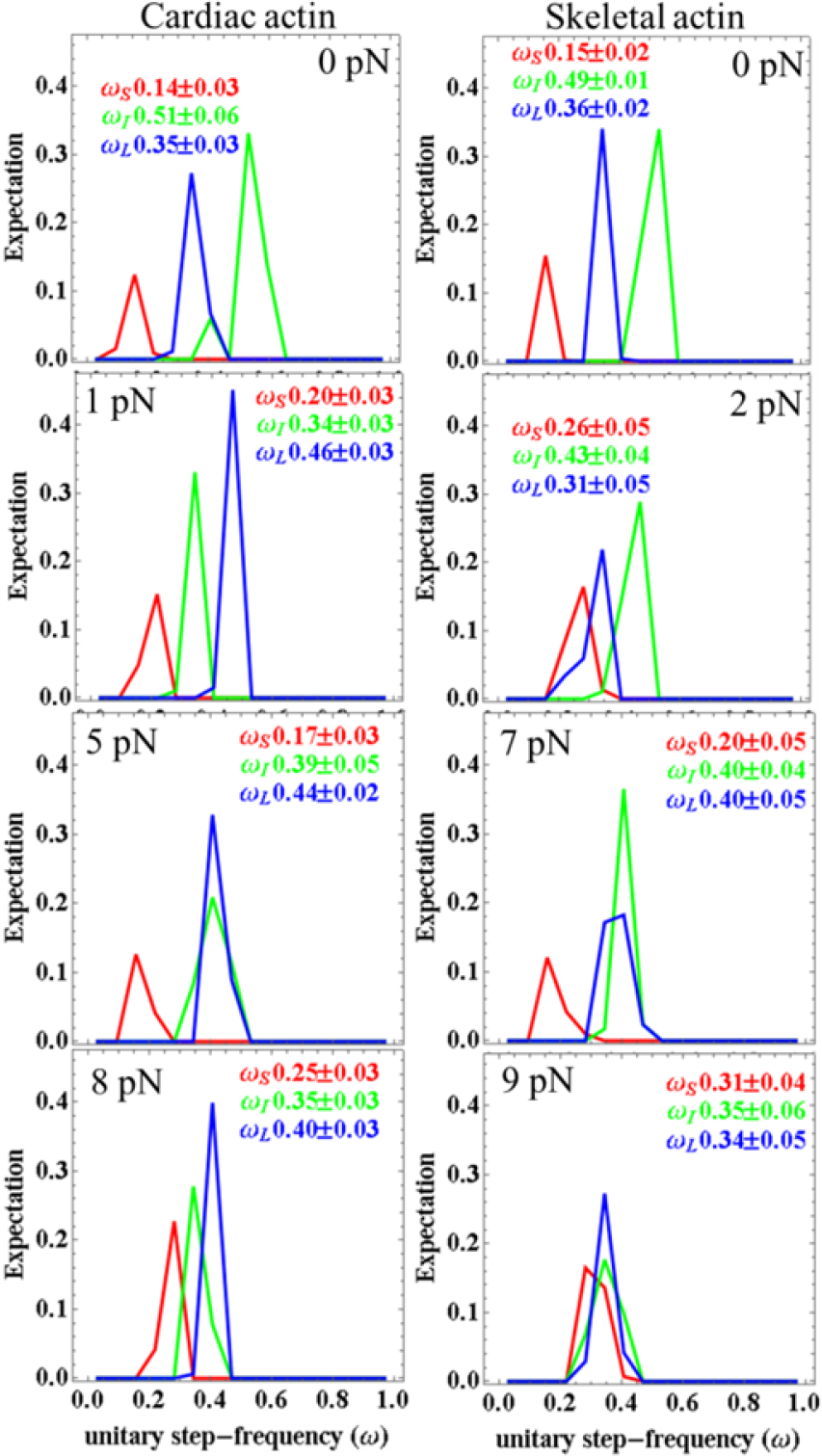
Step-frequencies for cardiac (left column) and skeletal (right column) actin with frictional loading indicated from 0 to 9 pN. Curves are from simulation of the corresponding event-velocity histograms in Figure 5 and as described in Methods. Errors are standard deviations for the 8-16 replicates indicated for each species in Figure 5. Leading step-size choice for βmys moving cardiac actin is always the long (blue) step-size. In contrast, βmys moving skeletal actin is always the intermediate (green) step-size. These selections reflect competitive regulation of the myosin mechanical characteristics by the ELC ratchet strain and lever arm strain mechanisms.

**Figure 7 panel a** shows average unitary displacement in nm, <d> in eq. 11, and **panel b** the duty ratio, <f> in eq. 12, for skeletal (red) and cardiac (blue) actin derived from simulation of each event-velocity histogram acquisition. Error bars show standard deviation for N acquisitions.

**Figure 7.**
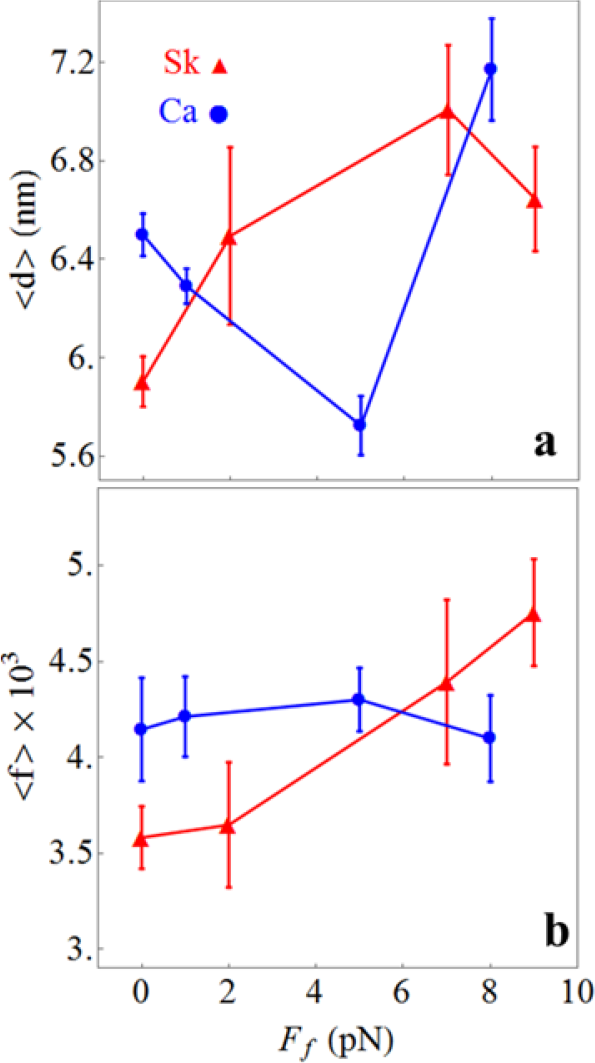
In vitro single βmys mean step-size <d> (panel a) and mean duty ratio <f> (panel b), for skeletal (red) and cardiac (blue) actin substrates measured with the Qdot assay under increasing load. Data points are derived from simulation of the corresponding event-velocity histograms in Figure 5 and as described in Methods. Errors are standard deviation for 8-16 replicates. Two-way ANOVA tested significance for factor 1 proteins skeletal actin and cardiac actin, and, factor 2 the frictional force F_f_ shows that <d> does not while <f> does differ significantly with confidence level p < 0.01. Nevertheless, <d> appears to signal the drag force-delayed onset of the high-force regime for myosin impelling cardiac actin since <d> increases dramatically to its skeletal value at the highest drag force.

Under unloaded conditions (top row in **Figures 5** & **6**) unitary step-sizes indicate a significant 0.4-0.5 nm shift to longer displacements for the cardiac actin while step-frequencies are identical for the two actins. Average displacement (**Figure 7 panel a**) also reflects a longer average unitary displacement for the unloaded (F_f_ = 0) cardiac actin. The different characteristic displacement is reflected equally in each unitary step-size implying a geometrical difference in the actomyosin complex probably unrelated to individual pathways in **Figure 2**. It implies that the overall 3-D structure of cardiac F-actin probably differs from skeletal F-actin giving the lever arm swing a longer projection in the direction of F-actin movement.

Although 3 distinct unitary step-sizes characterize the Qdot labeled actin motility in simulation, we now enjoy higher spatial resolution and observe a more complex pattern dependent on load where new step-sizes emerge and others decline demonstrating changing flux through alternative pathways in **Figure 2**. At or below 5 pN resisting force, the cardiac actin event-velocity histogram maintains the familiar short, intermediate and long step-sizes of ~3, ~5, and ~8 nm attributed to 5+3 nm (green-yellow), 5 nm (green), and 8 nm (blue) pathways in **Figure 2** but not including the solo-3^+^ nm pathway (red) that contributes at higher resisting force *^25^*. Low-force unitary step-sizes slightly shorten as resisting force increases as expected. This is the low-force regime for cardiac actin. At 5 pN resisting force, a nascent 4 nm step-size emerges in the cardiac actin event-velocity histogram (red asterisk, **Figure 5**). It is established in the event-velocity histogram by 8 pN (**Figure 5**) signaling onset of the high force regime for cardiac actin. During the transition to high force in the short step-size category the canonical 3 nm step-size declines in favor of an emergent new ~4 nm step-size. Step-size does not lengthen under resisting force but rather shortens as observed for the cardiac actin in the low-force regime and elsewhere *^58^* hence the new 4 nm step-size is from: the solo-3^+^ nm red pathway, a strain-shortened 5 nm step-size from the green pathway, or contributions from both (**Figure 2**). Skeletal actin data transitions to the high-force regime at or before the lowest resisting force of 2 pN.

The solo-3^+^ nm pathway produces a 4 nm step-size when slip distance is ~4 nm well within distance ranges previously reported *^44, 45^*. Disappearance of the ~3 nm step-size (from 5+3 nm yellow pathway) also signals reduction in the 5 nm step-size (from 5 nm green pathway), especially since lever arm strain inhibited ADP release would tend to boost 3 nm step-frequency. This observation is coupled with emergence of the ~4 nm step-size implying flux normally flowing into the green pathway at low force preemptively diverts to the solo-3^+^ nm (red) pathway and suggesting the ~4 nm step-size has minimal or no contributions from a strain-shortened 5 nm step-size. Thus the solo-3^+^ nm pathway solely causes emergence of the ~4 nm step-size at resisting forces >5 pN for cardiac actin and at >0 and <2 pN for skeletal actin.

In the high force regime, the intermediate step-size category has a characteristic displacement slightly rising to ~6 nm suggesting residual 5 nm step-size displacements combine with strain-shortened 8 nm step-size displacements from the blue pathway. The long step-size category shows displacement at 9-11 nm about equal to the sum of steps from the short and intermediate categories. The latter is unlikely to be unitary but rather separate unitary events from the short and intermediate categories sometimes falling into one frame capture interval and observed as a single peak in the event-velocity histogram.

Step-frequencies in **Figure 6** indicate cardiac actin shifts the leading step-frequency to the long step-size with application of the lowest resisting force. This was not observed previously and unique for the cardiac actin substrate. We suggest below that it is because cardiac actin stabilizes the ELC N-terminus binding by an extra hydrogen bond at Ser358 that is absent for Thr358 in skeletal actin. The short step-size frequency increases with loading force except at the transition from low- to high-force regimes in cardiac actin (≳5 pN resisting force) where the low-force short step-size probability is phasing out in favor of the high-force short step-size probability. The 3 step-size simulation inadequately quantitates flux through the system pathways during transition at 5 pN (**Figure 5**). Skeletal actin is always in the high-force regime (except at 0 load) where assigning density to unitary steps is ambiguous as discussed above hence skeletal actin step-frequencies in **Figure 6** are qualitative.

Duty-ratios in **Figure 7b** (<f>) compare cardiac and skeletal actin isoforms. Skeletal actin is always in the high force regime (except at 0 load) where assigning density to unitary steps is ambiguous as mentioned above hence quantities assigned to skeletal actin in **Figure 7** are qualitative. The cardiac actin isoform has constant or decreasing <f> over the frictional loading forces tested while skeletal actin has increasing <f>. The former is remarkable because it implies that the increasing load does not raise t_on_ as would be expected if lever arm strain-inhibited ADP release was the sole regulatory mechanism adjusting force-velocity to match load. Instead the system utilizes actively competing mechanisms of lever arm strain-inhibited ADP release and the ratcheting ELC N-terminus for constant or decreasing <f> by step-size downshifting. It contrasts with skeletal actin and suggests their difference must lie in the different ELC N-terminus actin binding affinities implying the weaker affinity skeletal actin interaction distorts competition between lever arm strain-inhibited ADP release and ratcheting ELC N-terminus mechanisms. A constant or decreasing duty-ratio may be a desirable characteristic for normal cardiac function by enforcing a constant or lower fraction of strongly bound myosins over most or all of the normal contraction cycle.

### Structure of the ELC/actin interaction in cardiac muscle

**Figure 8** shows ventricular myosin bound to cardiac actin at the end of the powerstroke based on a skeletal protein structure in the equivalent configuration from Aydt et al. *^18^*. The figure shows the 3 actin monomers interacting with one myosin S1 and including the N-terminus ELC/actin interaction. The four cardiac actin residues differing from skeletal actin (Asp2, Glu3, Leu299, and Ser358) are depicted with space filling atoms. Homology modelling indicates a potential hydrogen bond between sidechains Ser358 O-H and Glu6 O on the βmys ELC N-terminus actin binding domain. This interaction, and its homologue from all skeletal actomyosin, is shown in **Figure 9**. The proposed hydrogen bond in the cardiac actomyosin is indicated with the blue dashed line and the H–O distance of 1.65 Å. In the skeletal actomyosin model the closest approach of sidechain heavy atoms from Thr358 (skeletal actin) and Val6 (skeletal ELC) is 3.9 Å.

**Figure 8.**
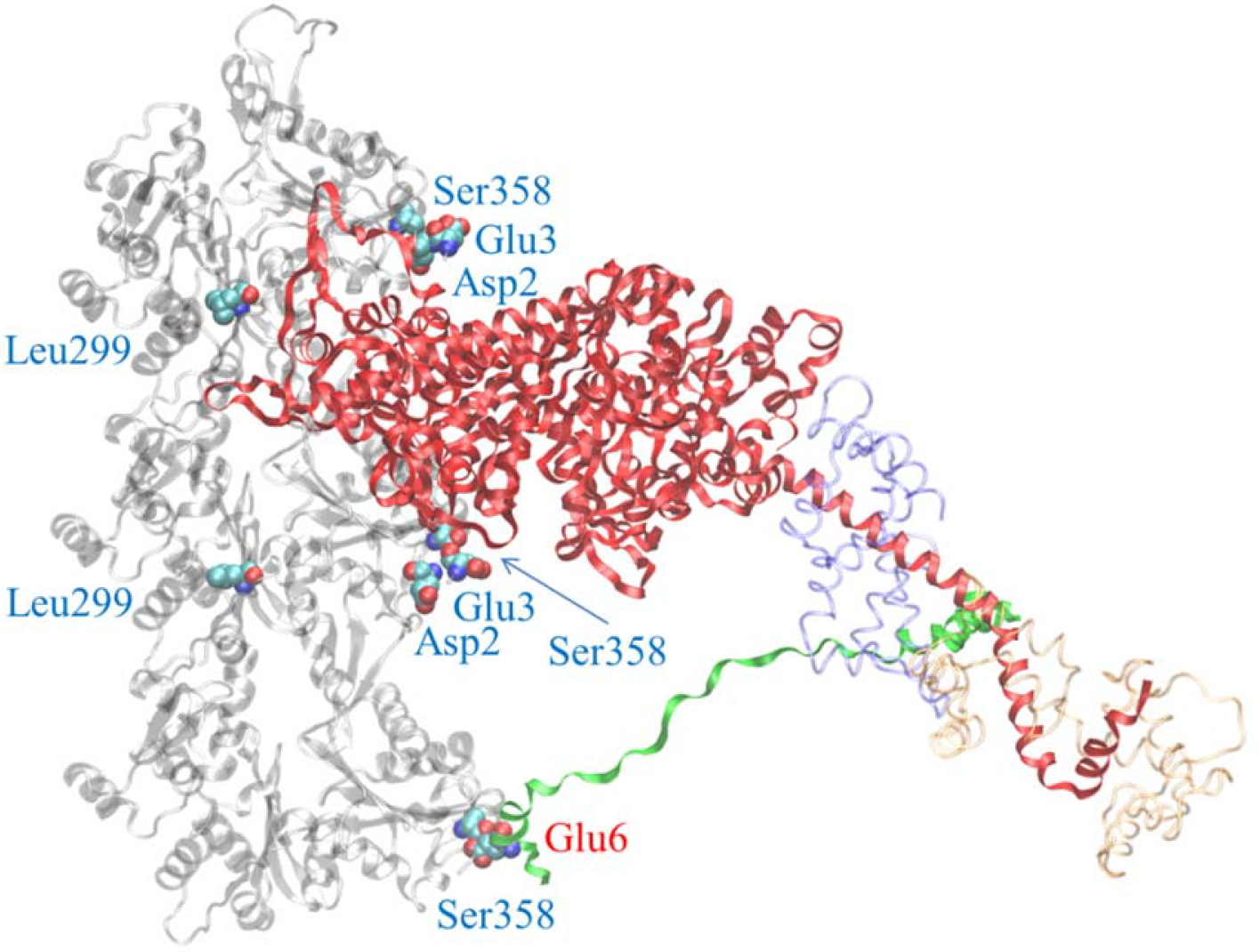
Homology modeled human ventricular myosin bound to cardiac actin and making the ELC/actin contact at the end of the powerstroke. Three cardiac actin crystal structures in a filament in transparent gray have the top two actin monomers contacting with the myosin heavy chain (red) and the bottom actin monomer in contact with the myosin ELC N-terminus (green). Myosin RLC (transparent red) and ELC not including the N-terminus (transparent blue) are indicated. Residues in the cardiac actin sequence differing from the skeletal sequence are denoted in space filling atoms with blue labels. Ser358 in the bottom actin monomer is proposed to form a hydrogen bond with one of the ∊-oxygens in Glu6 (denoted in space filling atoms with red label) from the ELC N-terminus (see Figure 9).

**Figure 9.**
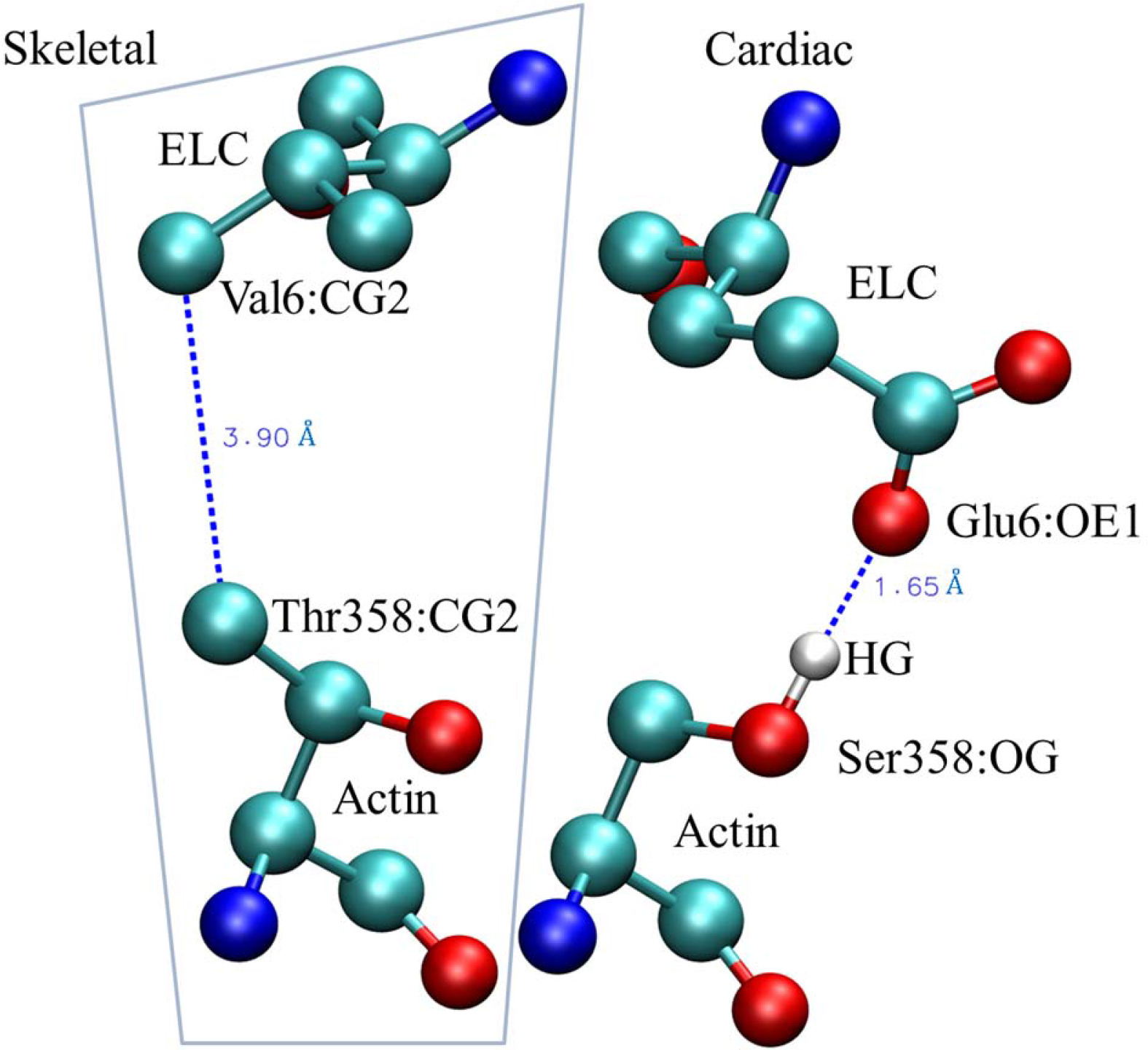
The actin Ser358 interaction with the ELC Glu6 from the cardiac isoforms (right) and the equivalent interaction for the skeletal actin and myosin sequences (left with boundary). The cardiac isoforms are proposed to enter into an O–H hydrogen bond depicted with the blue dashed line for atoms separated by 1.65 Å.

Removing cardiac actin contaminant from tissue purified porcine cardiac myosin is well known to be less efficient than removing the skeletal actin contaminant from tissue purified rabbit skeletal myosin *^4^*. We hypothesize that the actin/ELC-N-terminus contact for the cardiac actin and myosin isoforms stabilizes the cardiac actomyosin complex with the additional hydrogen bond between sidechains Ser358 O-H and Glu6 O (**Figure 9**) that is likely insensitive to ATP binding hence tending to frustrate efficient separation of cardiac actin from cardiac myosin in the presence of ATP.

## DISCUSSION

Skeletal and cardiac mammalian α-actin isoforms dominate in skeletal and cardiac muscle, respectively. They differ slightly from each other at 4 side chains where substitutions Glu2Asp, Asp3Glu, Leu299Met, and Thr358Ser convert skeletal to cardiac actin. In normal adult humans, skeletal muscle is homogeneous in the skeletal α-actin *^53^* and cardiac muscle has a skeletal:cardiac α-actin stoichiometry of ~1:4 *^29^*. Sequence similarity between isoforms promises near identical three-dimensional structures, nevertheless, they are functionally distinguishable with relative upregulation of the skeletal version in the ventriculum accompanying hypertrophic cardiomyopathy (HCM) disease *^30^*. We compared βmys ensemble enzymatic, ensemble mechanical, and single myosin mechanical characteristics when interacting with actin purified from skeletal and cardiac muscle. We find differences in the βmys ensemble performance due to the different actin substrates that we linked to a molecular mechanism using single myosin mechanical characteristics measured with the Qdot assay.

Actin-activated myosin ATPase V_max_ and K_m_ (**Figure 3** & **Table 1**) are statistically indistinguishable for the two actin isoforms indicating these substrates do not significantly affect motor domain ATP free energy transduction. It suggests the skeletal/cardiac actin substitutions Glu2Asp and Asp3Glu negligibly impact myosin functionality. In contrast, **Figure 4** compares the ensemble βmys force-velocity characteristics for skeletal and cardiac actin filaments. The Moore-Hill characterization *^38, 39^* specialized for this application indicates significant differences in ensemble motility velocity (**panel a**), force-velocity (**panel b**) and force-power (**panel c**) that unequivocally demonstrate functional differences between the two actin substrates. It is apparent from the ensemble data that skeletal actin upregulates power and consequently why dynamic heart tissue upregulates the skeletal actin substrate when heart disease compromises power production. We demonstrated using the Qdot assay that the molecular basis for the functional differences between by the two actin substrates lies in the strain-sensitive myosin regulation mechanisms represented in **Figure 2**.

Single myosin mechanical characteristics affected variously by the different actin isoforms are summarized in **Figures 5–6**. Unitary displacement classification constraints introduced here better separate signal from background giving higher step-size resolution for distinguishing 3 from 4 nm step-sizes. Emergence of the short step-size at ~4 nm contributed by the solo-3^+^ nm pathway (red in **Figure 2**) marks the onset of the high-force regime for both actins. For cardiac actin, the high-force regime begins to impact myosin function when the in vitro muscle (one actin filament moving over myosin in the assay) encounters ≳ 5 pN resisting force (**Figure 5**). Peak power for this isoform occurs for ~8 pN drag force. Skeletal actin in contrast reaches the high-force regime at >0 but ≲2 pN drag force (**Figure 5**). Uneven onset of the high-force regime for skeletal and cardiac actin by application of the resisting force implies dynamic hybrid force-velocity characteristics will emerge as the skeletal/cardiac actin fractional content increases in the disease compromised cardiac muscle. The emergent muscle will tend to activate the solo-3^+^ pathway at lower resisting force and with an upswing in sliding velocity, duty-ratio, and power.

A more subtle distinction between the actin isoforms is characterized in **Figure 7b** with average duty ratio, <f>. It shows the cardiac actin system utilizes actively competing mechanisms for lever arm strain-inhibited ADP release and the ELC ratchet to hold <f> constant. It contrasts with skeletal actin and suggests their difference must lie in the different ELC N-terminus actin binding affinities implying the weaker affinity skeletal actin interaction distorts competition between the lever arm strain mechanisms. A constant duty-ratio over dynamic loading may be a desirable characteristic for normal cardiac function by enforcing a constant fraction of strongly bound myosins over the normal contraction cycle in contrast to a modulating fraction presumably interfering with super-relaxation maintenance *^59, 60^*. The novel force-velocity characteristics for βmys caused by the actin isoforms in the heart muscle, and that are progressively modulated with heart disease onset, are dependent mainly on the cardiac myosin ELC and its weaker interaction with the skeletal actin at Thr358 (**Figure 9**). This insight, provided by the Qdot assay, elucidates the fundamental regulatory mechanisms in an autonomous myosin and identifies the site on actin responsible for the functional differences imposed by skeletal and cardiac actin.

In the absence of load, Qdot assay measured average and individual βmys step-sizes are 0.4–0.5 nm longer for the cardiac vs skeletal actin substrates (top row of **Figure 5**). Contacts between βmys heavy chain and the actin isoforms are identical based on actin-activated myosin ATPase hence it follows that differences between structures of the skeletal and cardiac actin rather than pathway flux regulation is responsible. Furthermore, residues differing between the isoforms do not fall within the contacts between G-actin monomers in F-actin *^61, 62^* implying a global F-actin structural difference causes the longer unitary step-sizes for βmys bound to cardiac actin. The βmys lever-arm swing when myosin makes the powerstroke causes greater displacement of the cardiac vs skeletal actin implying a greater nm per ATP efficiency under low loading for the heart. A minor efficiency improvement is a meaningful adaptation in a muscle that works constantly like the heart. Nonetheless, in conditions where energy conservation would seem critical, such as in fatigue resistant skeletal muscle, βmys is present but not cardiac actin *^63^*.

## CONCLUSION

Ensemble βmys force-velocity characteristics for skeletal and cardiac actin filaments compared in the Moore-Hill characterization (**Figure 4**) unequivocally demonstrate functional differences between the actin substrates. We investigated the molecular mechanism responsible for the differences by measuring single βmys step-size and step-frequency characteristics when moving loaded cardiac and skeletal actin using the Qdot assay. New Qdot assay data classification constraints introduced here better separates signal from background giving higher signal-to-noise data and permitting sub-nanometer step-size resolution. With this new technology we demonstrated significant myosin step-size distribution differences for skeletal vs cardiac actin substrate under loaded conditions.

We find that the skeletal and cardiac actin isoforms have different set-points in their strain-sensitive regulation such that onset of the high-force regime, where an increased contribution from the solo-3^+^ pathway (**Figure 2**) contributes a new 4 nm step-size displacement, is offset to higher loads when myosin drives cardiac actin. We attribute cardiac actin impact on myosin load sensitivity to the unique and specific cardiac ELC N-terminus/cardiac-actin contact at Glu6/Ser358 (**Figure 9**). It modifies the cardiac motor force-velocity characteristics detected with single myosin mechanics by stabilizing the ELC N-terminus/cardiac-actin association. Higher affinity at this site frustrates efficient actomyosin dissociation by ATP during βmys purification leading to more actin contamination (**Figure 1**) and is observed functionally to enhance long (8 nm) and short (3 nm) step-size probabilities at the expense of the intermediate (5 nm) step-size probability in the low-force regime (**Figure 6** for loads ≲ 5 pN). These observations combined with the constant duty ratio (**Figure 7 panel b**) show the cardiac actin system utilizes actively competing mechanisms of lever arm strain-inhibited ADP release and the ELC ratchet for maintaining approximately constant <f> over increasing load by step-size downshifting. In contrast, the skeletal actin substrate drives the system into the high-force regime at much lower loads to upregulate the solo-3^+^ nm pathway flux, power, and <f> (**Figures 4 and 7 panel b**). The latter characteristics are progressively upmodulated with HCM disease onset.

The 3-dimensional cardiac F-actin structure was shown to be modified compared to skeletal F-actin conformation. The structural change imparts a subtle but significant 0.4-0.5 nm myosin displacement advantage to the cardiac F-actin in vitro. It implies a greater efficiency for actin displacement vs ATP consumed.

## DATA ACCESSIBILITY

Data used in this study is provided in summary form in the text. Representative raw Qdot assay data is deposited at Zenodo (md5:86709883e2fc643bf67a61c84e66b2e0).

## ACKNOWLEDGEMENT

We thank Susanna P. Garamszegi for excellent technical assistance with preparation and characterization of the cardiac actin. This work was supported by the Mayo Foundation.

## REFERENCES

[1] Al-Khayat, H. A., Kensler, R. W., Squire, J. M., Marston, S. B., and Morris, E. P. (2013) Atomic model of the human cardiac muscle myosin filament, Proc. Natl. Acad. Sci. USA 110, 318–323.

[2] Huxley, H. E. (1969) The mechanism of muscular contraction, Science 164, 1356–1366.

[3] Rayment, I., and Holden, H. M. (1993) Myosin subfragment-1: structure and function of a molecular motor, Curr. Opin. Struct. Biol. 1993, 944–952.

[4] Ajtai, K., Garamszegi, S. P., Park, S., Velazquez Dones, A. L., and Burghardt, T. P. (2001) Structural characterization of β-cardiac myosin subfragment 1 in solution, Biochemistry 40, 12078–12093.

[5] Ajtai, K., Garamszegi, S. P., Watanabe, S., Ikebe, M., and Burghardt, T. P. (2004) The myosin cardiac loop participates functionally in the actomyosin interaction, J. Biol. Chem. 279, 23415–23421.

[6] Mornet, D., Pantel, P., Audemard, E., and Kassab, R. (1979) The limited tryptic cleavage of chymotryptic S-1:An approach to the characterization of the actin site in myosin heads, Biochem. Biophys. Res. Commun. 89, 925–932.

[7] Mornet, D., Bertrand, R., Pantel, P., Audemard, E., and Kassab, R. (1981) Proteolytic approach to structure and function of actin recognition site in myosin heads, Biochemistry 20, 2110–2120.

[8] Sutoh, K. (1983) Mapping of actin-binding sites on the heavy chain of myosin subfragment 1, Biochemistry 22, 1579–1585.

[9] Lorenz, M., and Holmes, K. C. (2010) The actin-myosin interface, Proc. Nat. Acad. Sci. USA 107, 5.

[10] Lowey, S., Waller, G. S., and Trybus, K. M. (1993) Function of skeletal muscle myosin heavy and light chain isoforms by an in vitro motility assay, J. Biol. Chem. 268, 20414–20418.

[11] Sherwood, J. J., Waller, G. S., Warshaw, D. M., and Lowey, S. (2004) A point mutation in the regulatory light chain reduces the step size of skeletal muscle myosin, Proc. Natl. Acad. Sci. USA 101, 10973–10978.

[12] Pant, K., Watt, J., Greenberg, M., Jones, M., Szczesna-Cordary, D., and Moore, J. R. (2009) Removal of the cardiac myosin regulatory light chain increases isometric force production, The FASEB Journal 23, 3571–3580.

[13] Greenberg, M. J., Kazimierczak, K., Szczesna-Cordary, D., and Moore, J. R. (2010) Cardiomyopathy-linked myosin regulatory light chain mutations disrupt myosin strain-dependent biochemistry, Proc. Natl. Acad. Sci. USA 107, 17403–17408.

[14] Burghardt, T. P., Josephson, M. P., and Ajtai, K. (2011) Single myosin cross-bridge orientation in cardiac papillary muscle detects lever-arm shear strain in transduction, Biochemistry 50, 7809–7821.

[15] Burghardt, T. P., and Sikkink, L. A. (2013) Regulatory light chain mutants linked to heart disease modify the cardiac myosin lever-arm, Biochemistry 52, 1249–1259.

[16] Sutoh, K. (1982) Identification of myosin-binding sites on the actin sequence, Biochemistry 21, 3654–3661.

[17] Timson, D. J., Trayer, H. R., and Trayer, I. P. (1998) The N-terminus of A1-type myosin essential light chains binds actin and modulates myosin motor function, Eur. J. Biochem. 255, 654–662.

[18] Aydt, E. M., Wolff, G., and Morano, I. (2007) Molecular modeling of the myosin-S1(A1) isoform, J. Struct. Biol. 159, 158–163.

[19] Dominguez, R., and Holmes, K. C. (2011) Actin Structure and Function, Annual Review of Biophysics 40, 169–186.

[20] Bobroff, N. (1986) Position measurement with a resolution and noise-limited instrument, Rev. Sci. Instrum. 57, 1152–1157.

[21] Thompson, R. E., Larson, D. R., and Webb, W. W. (2002) Precise nanometer localization analysis for individual fluorescent probes, Biophys. J. 82, 2775–2783.

[22] Stout, A. L., and Axelrod, D. (1989) Evanescent field excitation of fluorescence by epi-illumination microscopy, Applied Optics 28, 5237–5242.

[23] Wang, Y., Ajtai, K., Kazmierczak, K., Szczesna-Cordary, D., and Burghardt, T. P. (2015) N-terminus of Cardiac Myosin Essential Light Chain Modulates Myosin Step-Size, Biochemistry 55, 186–198.

[24] Burghardt, T. P., Sun, X., Wang, Y., and Ajtai, K. (2016) In vitro and in vivo single myosin step-sizes in striated muscle, J. Muscle Res. Cell Motil., 1–15.

[25] Wang, Y., Yuan, C. C., Kazmierczak, K., Szczesna-Cordary, D., and Burghardt, T. P. (2018) Single Cardiac Ventricular Myosins are Autonomous Motors, Open Biology 8, 170240.

[26] Nyitrai, M., and Geeves, M. A. (2004) Adenosine diphosphate and strain sensitivity in myosin motors, Philos. Trans. R. Soc. Lond. B. Biol. Sci. 359, 1867–1877.

[27] Burghardt, T. P., Ajtai, K., Sun, X., Takubo, N., and Wang, Y. (2016) In Vivo Myosin Step-Size from Zebrafish Skeletal Muscle, Open Biology 6, 160075.

[28] Burghardt, T. P., Sun, X., Wang, Y., and Ajtai, K. (2017) Auxotonic to Isometric Contraction Transitioning in a Beating Heart Causes Myosin Step-Size to Down Shift, PLoS ONE 12, e0174690.

[29] Vandekerckhove, J., Bugaisky, G., and Buckingham, M. (1986) Simultaneous expression of skeletal muscle and heart actin proteins in various striated muscle tissues and cells, J. Biol. Chem. 261, 1838–1843.

[30] Lim, D. S., Roberts, R., and Marian, A. J. (2001) Expression profiling of cardiac genes in human hypertrophic cardiomyopathy: insight into the pathogenesis of phenotypes, J. Am. Coll. Cardiol. 38, 1175–1180.

[31] Pardee, J. D., and Spudich, J. A. (1982) Purification of muscle actin, Methods Enzymol. 85, 164–179.

[32] Bergen III, H. R., Ajtai, K., Burghardt, T. P., Nepomuceno, A. I., and Muddiman, D. C. (2003) Mass spectral determination of skeletal/cardiac actin isoform ratios in cardiac muscle, Rapid Commun. Mass Spectrom. 17, 1467–1471.

[33] Wang, Y., Ajtai, K., and Burghardt, T. P. (2013) Qdot labeled actin super-resolution motility assay measures low duty cycle muscle myosin step-size, Biochemistry 52, 1611–1621.

[34] Wang, Y., and Burghardt, T. P. (2017) In vitro actin motility velocity varies linearly with the number of myosin impellers, Arch. Biochem. Biophys. 618, 1–8.

[35] Fiske, C. H., and Subbarow, Y. (1925) The Colorimetric Determination of Phosphorus, J. Biol. Chem. 66, 375–400.

[36] Ruhnow, F., Zwicker, D., and Diez, S. (2011) Tracking single particles and elongated filaments with nanometer precision, Biophys. J. 100, 2820–2828.

[37] Henriques, R., Lelek, M., Fornasiero, E. F., Valtorta, F., Zimmer, C., and Mhlanga, M. M. (2010) QuickPALM: 3D real-time photoactivation nanoscopy image processing in ImageJ, Nature Methods 7, 339–340.

[38] Greenberg, M. J., and Moore, J. R. (2010) The molecular basis of frictional loads in the in vitro motility assay with applications to the study of the loaded mechanochemistry of molecular motors, Cytoskeleton (Hoboken, N.J.) 67, 273–285.

[39] Hill, A. V. (1938) The Heat of Shortening and the Dynamic Constants of Muscle Proc. R. Soc. Lond. B 126, 136–195.

[40] Seow, C. Y. (2013) Hill’s equation of muscle performance and its hidden insight on molecular mechanisms, J. Gen. Physiol. 142, 561–573.

[41] Sugiura, S., Kobayakawa, N., Fujita, H., Yamashita, H., Momomura, S., Chaen, S., Omata, M., and Sugi, H. (1998) Comparison of unitary displacements and forces between 2 cardiac myosin isoforms by the optical trap technique. Molecular basis for cardiac adaptation, Circ.Res. 82, 1029–1034.

[42] Wang, Y., Ajtai, K., and Burghardt, T. P. (2014) Ventricular myosin modifies in vitro step-size when phosphorylated, J. Mol. Cell. Cardiol. 72, 231–237.

[43] Palmiter, K. A., Tyska, M. J., Dupuis, D. E., Alpert, N. R., and Warshaw, D. M. (1999) Kinetic differences at the single molecule level account for the functional diversity of rabbit cardiac myosin isoforms, J. Physiol. 519, 669–678.

[44] Kaya, M., Tani, Y., Washio, T., Hisada, T., and Higuchi, H. (2017) Coordinated force generation of skeletal myosins in myofilaments through motor coupling, Nature Communications 8, 16036.

[45] Debold, E. P., Patlak, J. B., and Warshaw, D. M. (2005) Slip Sliding Away: Load-Dependence of Velocity Generated by Skeletal Muscle Myosin Molecules in the Laser Trap, Biophys. J. 89, L34–L36.

[46] Mijailovich, S. M., Kayser-Herold, O., Stojanovic, B., Nedic, D., Irving, T. C., and Geeves, M. A. (2016) Three-dimensional stochastic model of actin–myosin binding in the sarcomere lattice, J. Gen. Physiol. 148, 459–488.

[47] Rayment, I., Rypniewski, W. R., Schmidt-Base, K., Smith, R., Tomchick, D. R., Benning, M. M., Winkelmann, D. A., Wesenberg, G., and Holden, H. M. (1993) Three-dimensional structure of myosin subfragment-1: A molecular motor, Science 261, 50–58.

[48] Kabsch, W., Mannherz, H. G., Suck, D., Pai, E. F., and Holmes, K. C. (1990) Atomic structure of the actin:DNAse I complex, Nature 347, 37–44.

[49] Holmes, K. C., Popp, D., Gebhard, W., and Kabsch, W. (1990) Atomic model of the actin filament, Nature 347, 44–48.

[50] Rayment, I., Holden, H. M., Whittaker, M., Yohn, C. B., Lorenz, M., Holmes, K. C., and Milligan, R. A. (1993) Structure of the actin-myosin complex and its implications for muscle contraction, Science 261, 58–65.

[51] Chen, L. F., Winkler, H., Reedy, M. K., Reedy, M. C., and Taylor, K. A. (2002) Molecular modeling of averaged rigor crossbridges from tomograms of insect flight muscle, J. Struct. Biol. 138, 92–104.

[52] Marti-Renom, M. A., Stuart, A. C., Fiser, A., Sanchez, R., Melo, F., and Sali, A. (2000) Comparative protein structure modeling of genes and genomes, Annu.Rev.Biophys.Biomol.Struct. 29, 291–325.

[53] Ilkovski, B., Clement, S., Sewry, C., North, K. N., and Cooper, S. T. (2005) Defining α-skeletal and α-cardiac actin expression in human heart and skeletal muscle explains the absence of cardiac involvement in ACTA1 nemaline myopathy, Neuromuscul. Disord. 15, 829–835.

[54] Behrmann, E., Müller, M., Penczek, Pawel A., Mannherz, Hans G., Manstein, Dietmar J., and Raunser, S. (2012) Structure of the Rigor Actin-Tropomyosin-Myosin Complex, Cell 150, 327–338.

[55] Murphy, C. T., and Spudich, J. A. (1999) The sequence of the myosin 50-20K loop affects myosin’s affinity for actin throughout the actin-myosin ATPase cycle and its maximum ATPase activity, Biochemistry 38, 3785–3792.

[56] Várkuti, B. H., Yang, Z., Kintses, B., Erdélyi, P., Bárdos-Nagy, I., Kovács, A. L., Hári, P., Kellermayer, M., Vellai, T., and Málnási-Csizmadia, A. (2012) A novel actin binding site of myosin required for effective muscle contraction, Nat Struct Mol Biol 19, 299–306.

[57] Ranatunga, K. W. (1984) The force-velocity relation of rat fast- and slow-twitch muscles examined at different temperatures, J. Physiol. 351, 517–529.

[58] Kaya, M., and Higuchi, H. (2010) Nonlinear elasticity and an 8-nm working stroke of single myosin molecules in myofilaments, Science 329, 686–689.

[59] Hooijman, P., Stewart, M. A., and Cooke, R. (2011) A new state of cardiac myosin with very slow ATP turnover: A potential cardioprotective mechanism in the heart, Biophys. J. 100, 1969–1976.

[60] Alamo, L., Pinto, A., Sulbarán, G., Mavárez, J., and Padrón, R. (2017) Lessons from a tarantula: new insights into myosin interacting-heads motif evolution and its implications on disease, Biophysical Reviews.

[61] Sawaya, M. R., Kudryashov, D. S., Pashkov, I., Adisetiyo, H., Reisler, E., and Yeates, T. O. (2008) Multiple crystal structures of actin dimers and their implications for interactions in the actin filament, Acta Crystallogr. Sect. D. Biol. Crystallogr. 64, 454–465.

[62] von der Ecken, J., Müller, M., Lehman, W., Manstein, D. J., Penczek, P. A., and Raunser, S. (2014) Structure of the F-actin–tropomyosin complex, Nature 519, 114.

[63] Vandekerckhove, J., and Weber, K. (1979) The complete amino acid sequence of actins from bovine aorta, bovine heart, bovine fast skeletal muscle and rabbit slow skeletal muscle, Differentiation 14, 123–133.

